# 4D imaging and analysis of multicellular tumour spheroid cell migration and invasion

**DOI:** 10.1101/443648

**Authors:** Rosalie Richards, David Mason, Rebecca Kelly, Raphaël Lévy, Rachel Bearon, Violaine Sée

## Abstract

Studying and characterising tumour cell migration is critical for understanding disease progression and for assessing drug efficacy. Whilst tumour cell migration occurs fundamentally in 3 spatial dimensions (3D), for practical reasons, most migration studies to date have performed analysis in 2D. Here we imaged live multicellular tumour spheroids with lightsheet fluorescence microscopy to determine cellular migration and invasion in 3D over time (4D). We focused on glioblastoma, which are aggressive brain tumours, where cell invasion into the surrounding normal brain remains a major clinical challenge. We developed a workflow for analysing complex 3D cell movement, taking into account migration within the spheroid as well as invasion into the surrounding matrix. This provided metrics characterising cell motion, which we used to evaluate the efficacy of chemotherapeutics on invasion. These rich datasets open avenues for further studies on drug efficacy, microenvironment composition, as well as collective cell migration and metastatic potential.

## Introduction

The multicellular tumour spheroid is a well-characterised model in which cells are grown as three-dimensional (3D) clusters in the absence of any exogenous support. The spheroid model replicates important aspects of the tumour microenvironment such as gradients in oxygen, nutrients, pH, and cell proliferation and survival (1–3). The cellular and molecular heterogeneity of spheroids makes them a better model for solid tumours than homogeneous monolayer cultures. These pathophysiologically-representative 3D models are increasingly important in cancer research, especially for drug testing.

The invasion of tumour cells into neighbouring tissue or distant organs is a hallmark of cancer and a major cause of mortality (4), which urgently needs to be tackled therapeutically. Good *in vitro* models for testing drugs that inhibit cell invasion are therefore essential. The majority of in *vitro* invasion assays to date, and associated drug screening, have been conducted using cells cultured as two-dimensional (2D) monolayers on plastic and glass surfaces (5). These conditions fail to replicate important aspects of the cellular environment that occur *in vivo*, such as complex cell-cell and cell-matrix interactions, matrix stiffness, and gradients in soluble factors such as oxygen, nutrients, growth factors, and cytokines (6).

This explains why the results of *in vitro* drug screening often only poorly correlate with results obtained in vivo. Hence, drug testing in spheroids is emerging as a beneficial intermediate between testing in monolayer cultures and testing in animals (7). In previous attempts at measuring cellular invasion in 3D models, the spheroids were plated on, or embedded in, extracellular matrix (ECM) in 96-well plates and imaged using an inverted microscope (8–11) providing information about the movement of some of the invading cells (in the x-y plane) but failing to capture the movements of cells within the spheroids. Imaging thick, living samples over long time courses can be challenging due to the effects of light diffraction, photobleaching and phototoxicity (12).

Lightsheet fluorescence microscopy (LSFM), also known as single-plane illumination microscopy (SPIM), has overcome some of these challenges. By illuminating only the focal plane of the detection objective, the effects of photobleaching and phototoxicity are much reduced compared to confocal microscopy (13, 14). LSFM also allows for very fast image acquisition from multiple angles, making it the ideal technique for imaging cell movement in a large, tightly packed spheroid, where z-stacks need to be acquired in rapid succession to facilitate the tracking of cells in space. Concurrently, the development of new software tools for spatial-temporal quantification of tumour spheroid dynamics (15) facilitates the analysis of such data rich 3D time lapse experiments. Previous examples of live 3D spheroid imaging using LSFM include the monitoring of cell division and nuclear shape in 3D by imaging human colon carcinoma cells stably expressing a fluorescent H2B (16, 17).

Here we used live LFSM of multicellular tumour spheroids to characterise cellular migration and invasion in 4D. We developed a workflow for analysing complex 3D cell trajectories with either commercial or open source software platform for tracking, taking into account migration within the spheroid as well as invasion into the surrounding extracellular matrix. This generated metrics characteristic of cell motion. These methods are applicable to a range of experimental conditions, including anti-cancer drug treatment. As proof of principle, spheroids were treated with 17-allylamino-17-demthoxygeldanamycin (17AAG), a heat-shock protein 90 (HSP90) inhibitor, and cilengitide, an *α*v*β*3 and *α*v*β*5 integrin inhibitor with anti-tumour activity (18). Live LSFM combined with cell motion analysis allowed us to discern the effects of these molecules both on the cells invading away from the spheroid as well as on movement within the spheroid. Both effects may need to be considered to evaluate the mechanisms and efficacy of anticancer drugs in inhibiting tumour cell infiltration.

## Methods

### H2B-mRFP-U87MG stable cell line production

A nuclear marker is necessary to facilitate object recognition by the tracking software and enable automated cell tracking. We previously found that nuclear stains with dyes show poor penetration in spheroids, therefore, we created a cell line stably expressing a fluorescent histone H2B nuclear reporter protein by transducing U87-MG cells with pHIV-H2BmRFP. Lentiviral particles were generated by transfecting HEK293T cells with the following plasmids from Addgene (http://www.addgene.org): pHIV-H2BmRFP (Plasmid #18982), psPAX2 (Plasmid #12259) and pMD2.G (Plasmid #12260) in a 4:2:1 ratio. On day 1, HEK 293T cells were seeded in six 10 cm^2^ dishes at a density of 1.5 x 10^6^ per dish in 10 mL growth media (DMEM + 10% FBS + nonessential amino acids). On day 2, cells were transfected with the aforementioned plasmids using polyethylenimine (PEI) at a ratio of 2 *μ*L: 1 *μ*g PEI:DNA. On day 3, the media was changed and on day 5 supernatant was harvested, centrifuged at 500 x g for 10 min and filtered with a 0.45 *μ*m PES filter before being subjected to ultracentrifugation following the protocol described in (19). 2.5 x 10^5^ U87-MG cells were seeded in to a 25 cm^2^ tissue culture flask and 250 *μ*L virus was applied. Media was changed after 24 h.

### Spheroid culture and preparation

We have previously described a workflow for the growth, handling and imaging of spheroids made with fluorescently-labelled cells (20). The hanging drop method was used to grow compact spheroids from U87-H2BmRFP glioblastoma cells over 3 days (detailed protocol is available in (20)). Three day old spheroids were used to limit the effect of light scattering seen when imaging deeper (past ~150 *μ*m). At the beginning of imaging, spheroids were approximately 300 *μ*m in diameter. Spheroids were prepared for imaging by aspirating the growth media and replacing it with 50% ice-cold Matrigel (Corning, Netherland) and 50% growth media plus 25 mM HEPES. For matrigel labelling, 1:1000 dilution of 5 mg/ml FITC-dextran (Sigma, MW 148,000) was added in the growth media. For drug treated spheroids, 0.5 *μ*M 17AAG (Abcam, UK) or 2 *μ*M cilengitide (Adooq Bioscience, CA, USA) was added to the growth media. A Teflon plunger (Zeiss) was used to draw the spheroid into an optically-amenable fluorinated ethylene propylene (FEP) tube (S 1815-04, BOLA, Germany), which was sealed at one end using parafilm, mounted in the Lightsheet Z.1 (Zeiss) sample holder and then immersed in water at 37 °C for the duration of the experiment.

### Image acquisition

Samples were excited with a 561 nm laser and 10x excitation objectives; emitted light was collected through a 576–615 band pass filter using a 20x W Plan-Apochromat objective and 1x zoom. Images of the H2B-mRFP-labelled nuclei were acquired every 3 min with a z-step of 1.66 *μ*m, starting approximately 50 *μ*m in front of the spheroid. Lightsheet Z.1 Zen software (Zeiss) was used to acquire images using online dual side fusion with pivot scan. Images were detected using a pco.edge scientific complementary metal–oxide–semiconductor (sCMOS) camera.

### Image pre-processing

Processing was done on a high-end workstation (*cf*. a high performance compute cluster), and thus reducing the size of the lightsheet data prior to analysis was essential due to the large file sizes produced (up to 0.75 TB for one experiment). A custom Fiji macro (see code availability below) allowed us to load one time point at a time and to down-sample the data by decreasing bit depth from 16 bit to 8 bit and by performing 2x binning (this is user configurable). A further processing step was carried out in Fiji prior to analyses in order to deal with sample drift (21), using the descriptor-based series registration plugin developed by Preibisch et *al*. (22). Registration was carried out using a translational transformation model and consecutive matching of images with a range for all-to-all matching of two.

### Cell tracking

There are many excellent software packages available for tracking points over time (23). In this study, the data presented were tracked using the commercially-available Imaris 8.2.0 software (Bitplane, Belfast). The accuracy of the tracking (specifically the feature linkage) were assessed by selecting random tracks from each experiment and confirming that linkage was being correctly assigned. In some cases, imperfect linkages were identified due to changes in feature quality; however, our analysis does not rely upon having complete tracks.

The most basic tracking outputs are:

1. Position (Cartesian coordinates in X, Y and Z)
2. Time stamp (calibrated time or frame number)
3. Track ID to link features across frames

Although most software provides further analysis such as calculating overall track speed or intensity of spots over time, to give our analysis the widest utility, we wrote our processing scripts to assume no *a priori* track analysis information other than these five essential outputs. Tracking was carried out using an autoregressive motion algorithm, with a maximum distance of 6 *μ*m and a maximum gap size of zero. A filter was applied to remove any tracks that were shorter than eight time points (24 min). Once tracking was complete, the ‘Position’ spot statistics were exported into a CSV file. To make the analysis accessible to a wider range of researchers, the analysis script can also accept the ‘Spots in tracks statistics’ output from the open source Trackmate (24) plugin (v 3.8.0) for Fiji.

CSV outputs were imported into the MATLAB script for further processing.

### Cell division events

U87-H2BmRFP glioblastoma cells were seeded into wells of a 24-well plate then imaged on an epifluorescent microscope at 10x magnification in fluorescence and transmitted channels acquiring images every 150 seconds for 24 h. Time-lapse movies were registered using the descriptor-based series registration plugin (see ‘Image pre-processing’ above) using only frame-to-frame registration. Cells were tracked using Trackmate (24), applying a Linear Assignment Problem (LAP) feature linker which allows track split events to be identified. These are recorded as a proxy for cell divisions.

### Measurement of spheroid boundary and centroid

The spatial location of the outer boundary of each spheroid was defined by excluding the 10 outermost spots (to take into account cells released into the gel during loading) and taking the mean radial distance of the ten next furthest out spots present in the first frame of the experiment. A further 5% is added to the distance to take into account cell size. Because part of the analysis relies upon the isotropic nature of a spheroid, the coordinates of the spheroid centroid has to be provided by the user, which allows characterisation of the position and distribution of cells. This is requested from the user at the beginning of the data processing script. It is important to use the same software for estimating centroid position as for tracking, as different software may use different frames of reference to define the image origin. Using Imaris, three orthogonal clipping planes were created and positioned centrally to allow the read-out of the relevant position on each axis. The same result can be achieved in Fiji by measuring the X,Y & Z coordinates using the cursor position to read out on the status bar, or alternatively, using the centroid position from a segmented volume.

In Figure 1C, the density of the spheroids is calculated by converting the Cartesian (XYZ) coordinates into a spherical coordinate system (radial distance, azimuth and elevation). The axis of imaging corresponded to an elevation of zero degrees, thus cells were counted within a spherical section of 30 degrees elevation (see schematic in Figure 2Bi). The spherical section was separated into sampling bins by filtering only those spots whose radial distance fell between two consecutive boundaries 20 *μ*m (radial distance) apart out to 400 *μ*m (illustrated in 2D in Figure 1Bii) which was enough to encompass all spheroids in this study. The number of cells within this volume was recorded and then divided by the section volume to give the density in units of cells per *μ*m^3^.

**Figure 1.**
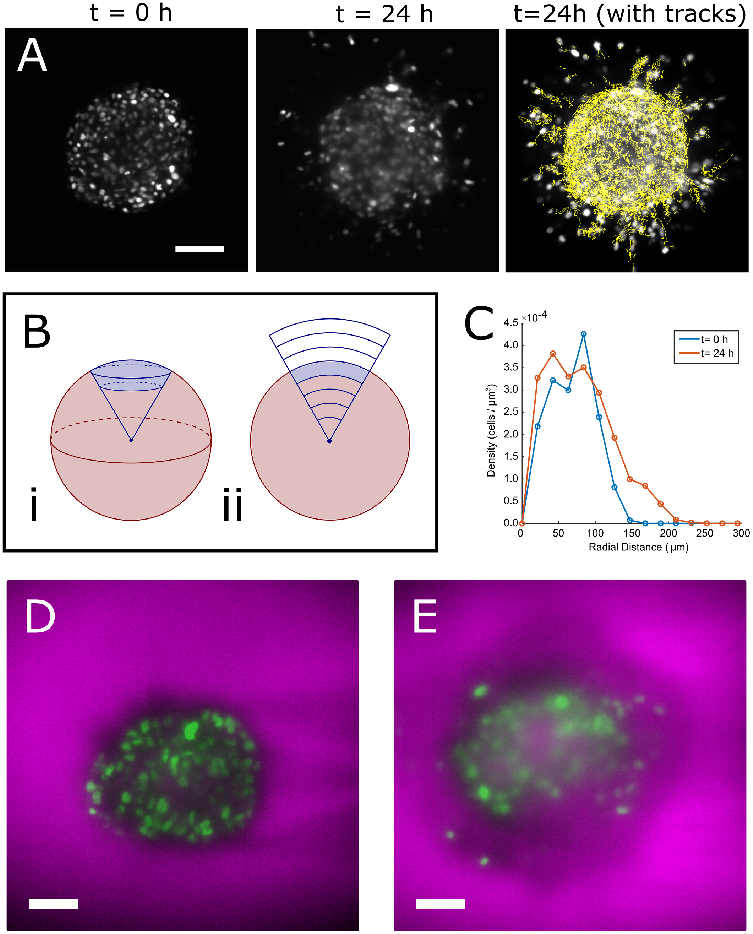
The spheroid as a model. Glioblastoma spheroids made with U87MG cells expressing H2B-mRFP were grown for 3 days, mounted on a lightsheet microscope (t=0 h) and imaged for up to 24 h. (**A**) Single slices of the 3D stack (left, middle). Invasion of individual cells can be tracked as they invade the surrounding matrix (tracking is done on full 4D datasets and tracks overlaid on the 24 h time-point in the rightmost panel). The scale bar represents 100 *μ*m. The density of the spheroid is calculated by counting the number of cells within each truncation of a spherical sector. This is shown in 3D in (**Bi**) and for clarity only, in 2D (**Bii**). The density (**C**) is described as the number of cells divided by the calculated volume of each truncation (see Methods). (**D-E**) The spheroid was embedded in a mix of FITC-dextran and Matrigel to demarcate the gel boundary (H2B-mRFP in green, dextran shown in magenta). The images show a rendering of a spheroid at t=0 h (**D**) and at t=24 h (**E**) Scale bars in D and E represent 50 *μ*m.

### Analysis of tracking data

MATLAB (Mathworks, US) was used to process the tracking output obtained from the analysis workflow (Supplemental Figure 1). A processing script was used to calculate spot and track parameters and a second script was used to create a characteristic set of plots. Briefly, an output CSV file was selected and, when requested, the Cartesian coordinates of the spheroid centroid and the time calibration (sec / frame) were entered. The boundary of the spheroid was calculated by excluding the outermost 10 features (we noted a few cells embedded in the gel during mounting) then taking the mean radial distance of the next 10 outermost features in the first frame of the movie and dilating this 5% to take into account spheroid drift (section 10 of the processing script allows for customisation of these parameters to fit individual setups). As spheroid drift was corrected in image pre-processing, we found this to be a reliable measure of the boundary (see Discussion section for commentary on suitability and also compare Figure 1D and 1E).

Without considering individual tracks the feature positions were used to measure the bulk changes in the spheroid over time by considering their radial position from the centroid. The cumulative distribution plots were created by counting the cumulative number of features identified for any radial distance from the centre. The figures reported in Supplementary Figure 5 are the result of a non-linear least squares fit of a generalised logistic function of the following form:

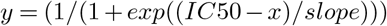

From spot features, the density can also be calculated (Figure 1C). When tracks are considered, the speed (3D interframe distance divided by interval), windowed speed (path length over a number of steps divided by the time taken) and straightness (calculated in a five frame window by dividing the end to end displacement over five frames by the sum of the displacements along the path length). The time-windowed approach (of 5 frames) was necessary to smooth the data while retaining good granularity. At track level, the mean squared displacement (MSD) was calculated with time lags up to 25% of the length of the individual track. We calculated the diffusion coefficient (D) using a linear regression to the first 10 points of each MSD/t plot, where the slope is equal to 6D for motion in three dimensions.

One of the key functionalities of the processing script is to break apart individual tracks into segments that are within the original spheroid boundary (defined at the start of the experiment) and those outside the boundary. The same statistics calculated for track-level data are provided as the mean of all like (IE inside or outside) segments.

### Plotting and visualisation

The main output of the processing script is a MATLAB workspace (`.mat`) file containing all of the calculated values for the features and tracks (see the code repository for a detailed description of the output tables). For meaningful visualisation, a second script was created, providing plots, including the summary plots shown in the figures. The plotting and output options of this script are controlled by a number of variables whose parameter space and functions are detailed in the script header (for example axis line widths, font sizes, etc). Other than the input and initialisation code section, each plot is written as a MATLAB section allowing the code to be run in completion or in sections to output only the desired figures.

### Statistical analysis

Data were processed to compare speed distribution. Statistical significance was determined using two-proportions z-test package in R (see https://www.rdocumentation.org/packages/stats/versions/3.6.0/topics/prop.test for more details).

### Code Availability

The pre-processing Fiji script and the processing and visualisation MATLAB scripts are available at https://bitbucket.org/davemason/lsfm_scripts along with documentation and example tracking data. At time of publication the commit hash is 9958866.

## Results

### The spheroid as a model system

Previous attempts to describe the migration of spheroid-associated cells over time have largely focussed on gross morphological changes to the spheroid itself, using 2D imaging or 3D projections (25). Such depictions may miss important dynamics occurring at a single cell level or within the body of the spheroid. This is shown in Figure 1A, where a single 2D section of two time points is displayed. To characterise single-cell movement, we developed an analysis pipeline (Supplemental Figure 1) to quantify the migration of single cells through postacquisition tracking and track analysis (see overlaid tracks in Figure 1A).

When investigating the motion of cells within a spheroid and into the surrounding matrix, an important consideration is the location of the boundary. Using the positions of the nuclei in all time points, we calculated the density of the spheroids throughout a spherical sector (shown in Figure 1Bi, clarified for a 2D case in 1Bii). Predictably, the density decreased at the estimated radius of the spheroid (Figure 1C, blue line). The low density at radial distances below ~50 *μ*m, is not due to a lack of cells or a necrotic core (which is not typically seen in spheroids this size) but to the difficulty imaging several hundred microns into an optically dense structure, hence reducing the number of identified features at this depth. The density towards the edge of the spheroid does drop as expected. Density measurement at the end of a 24 h time course, shows some expansion at the edge of the spheroid compared to time 0 h (Figure 1C, orange line). However, the core does not appear to enlarge (see Figure 1A and Supplementary Movie 1), as suggested by the change in density. To better visualise the boundary, we labelled the surrounding matrix by spiking the Matrigel with a high molecular weight fluorescent dextran. Over the course of ~18 h there was only a small increase in the measured diameter of the void occupied by the spheroid (from 160 to 178 *μ*m, Figure 1D, E). Given the relatively small change in void size over the course of the experiment, we decided to calculate the boundary based upon the position of the cells furthest away from the spheroid centroid in frame 1 after excluding possible outliers. We found this to be the most robust way to define the boundary in this dataset.

### Single track analysis in 3 dimensions

In the control spheroids, we observed cell migration within the spheroid as well as invasion into the surrounding matrix (Supplemental Movie 1). We tracked cell movements and obtained 47,771 tracks from 3 spheroids. One such track, showing the movement of a single cell, is shown in Figure 2A. Yet, the challenge is to extract parameters that allow the behaviour of the population of cells as a whole to be captured, in particular in the context of invasion. It is therefore critical to assess the distribution of single track or single spot data. Given that cells undergo migration within the spheroid, we hypothesised that speed (Figure 2B) and straightness (Figure 2C) could be informative metrics for describing the motion of cells. Both increased speed and straightness (the latter being a measure of the tortuosity of a track) could increase the likelihood of cells migrating in such a way to move outside the bounds of the original spheroid into the surrounding matrix.

**Figure 2.**
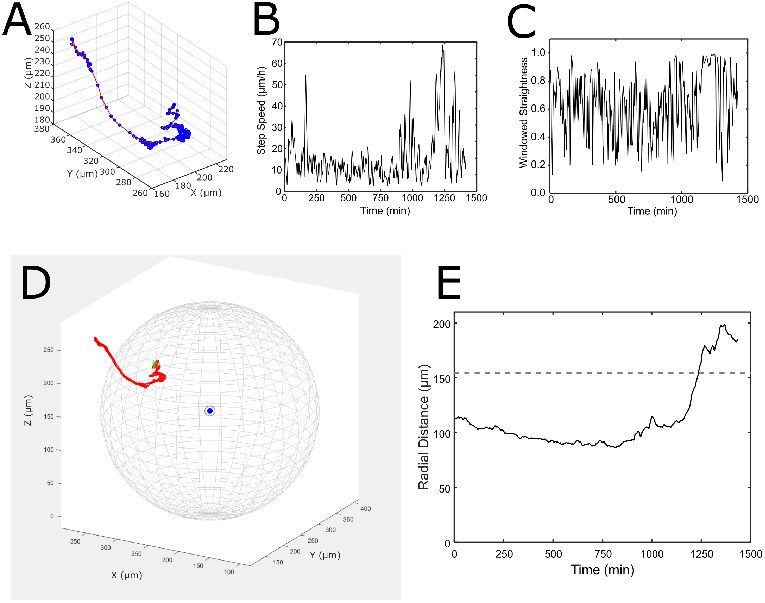
Single Trajectory Analysis. (**A**) A single track from a control spheroid is shown plotted in 3D. (**B**) Step speed and (**C**) straightness (displacement divided by path length over 5 frames) plotted over time. (**D**) The position of the track from its start point (green dot) in relation to the boundary can be visualised, given the coordinates of the spheroid centroid (blue dot). (**E**) The radial distance is used to record migratory events (dashed line represents the spheroid boundary).

Unlike track analysis in an embryo (26) or tissues, which can be quite amorphous, spheroids have roughly the same spatial properties in all directions with regards to the centre. With a defined centroid, a boundary can be demarcated between the edge of the spheroid and the surrounding matrix (see Methods) and thus all tracks have a geometric frame of reference (Figure 2D). The spot and track positions can therefore be characterised by their radial distances, and sorted into categories depending upon whether they remain within the spheroid boundary or cross the boundary and invade the surrounding matrix during the course of the experiment (see Figure 2E).

### Cell Population Analysis

One of the most basic features of metastatic cancer cells is invasion; i.e. their ability to leave their initial niche and permeate the nearby microenvironment. However, there is no clear metric to define and characterise invasion and the extent of invasion is often underestimated through the projection of 3D+time experiments onto 2D+time, either through imaging (for example by imaging a single thick z-plane) or post-acquisition using a projection in the z-axis. By locating cell nuclei in 3D space, we captured an accurate picture of invasion into the surrounding matrix and represented this through a cumulative distribution plot (Figure 3A). This type of plot describes the location of the cells at the start of the experiment (blue line shows frame 1) and how the distribution of the cells’ radial positions changes over the course of the experiment. Compared to a 2D projection, one important advantage is that information in the z-plane is preserved. Two interesting insights are gleaned from this plot. Firstly, the increase in the plateau (Figure 3A) in the raw data plots indicates an increased number of features being located at later time points. This can be explained by 1) cell division, clearly observable throughout the time course movies and 2) more cells being detected as they move into a higher contrast area of imaging. Secondly, if the plots are normalised (Figure 3B), the shift of the curve to the right, and indeed the shape of the curve itself, indicates changes in the distribution of the cells over time. We were unable to conclude whether cells were primarily moving from the edges of the spheroid into the matrix or whether there was a global expansion across the whole spheroid, yet the dextran labelling of the matrix suggested that there was little spheroid expansion (Figure 1D,E).

**Figure 3.**
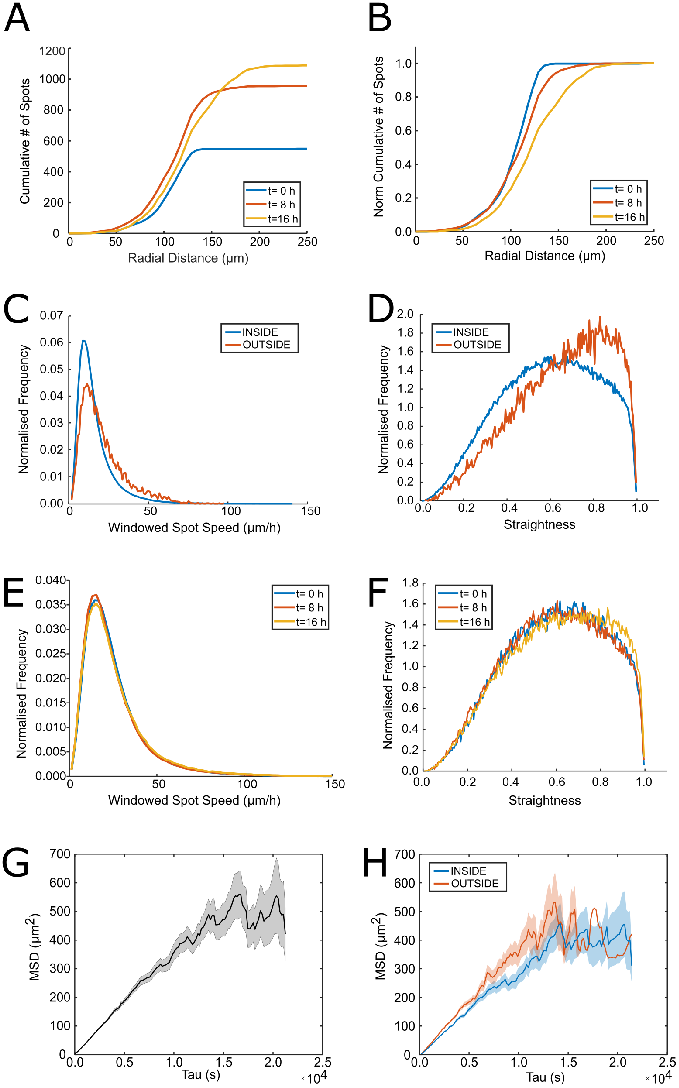
Population Level Analysis. (**A**) The cumulative distribution of detected nuclei by radial distance from the centroid is shown at three different time-points for a control spheroid. (**B**) The graph shown in (A) is normalised for each curve such that the plots plateau at a value of 1 to more easily observe the changing shape of the distribution. (**C,E**) Distribution of windowed spot speeds (over 5 frames) and (**D,F**) straightness (over 5 frames). Data shown as a function of radial distance (**C,D**) in two bins; inside (blue line; in this spheroid, a radial distance of 0-170 *μ*m) and outside (orange line; a radial distance greater than 170 *μ*m). (**C**) The proportion of fast speed observations from cells categorized as ‘outside’ (speed >20) is significantly greater than the proportion of fast speed observations from cells categorized as ‘inside’ with p-value < 2.2 x 10^-16^. Data shown as a function of experiment time (**E,F**) in three bins each representing the following 8 hours. Plots C-F are normalised to have equal area. (**G**) The Mean Square Displacement (MSD) was calculated (see Methods for details) for whole tracks. (**H**) The MSD was calculated for segments of tracks inside or outside the spheroid. Shaded areas represent the SEM, the plots represent 11255 segments inside the spheroid and 2493 outside.

The cumulative distribution of features effectively ignores the movement of cells in favour of their position at a given time; however, the tracking data are incorporated in the analysis and thus the distribution of speed and straightness parameters can be studied both as a function of radial distance (Figure 3C, D) and as a function of time (Figure 3E, F). We can show, for example, a mean track speed (± standard deviation) of 27 ± 10 *μ*m/h within the spheroid boundary (blue line Figure 3C) and 29 ± 12 *μ*m/h in the surrounding matrix. Analysing migration as a function of time and spatial localisation can provide useful insights. For example, drug treatment may effect cell speed and location and these effects may be further dependent on cell density, location within the spheroid, cell-cell contacts, drug diffusion / half-life, or activation of signalling feedback loops etc. Here, distance is especially relevant given the composition and physical properties of the embedding gel. The normalised plots in Figure 3C, D are shown with two bins of spatial data corresponding to tracks inside vs outside (based on the initial size of thespheroid - see details in Methods).

We initially calculated spot speed at the level of single steps, by taking the single-frame displacement of spots and dividing it by the acquisition interval (as in Figure 2B). This method provided noisy results, presumably due to the erratic motion often seen in migrating cells. To overcome this problem, we calculated windowed parameters, which have the effect of smoothing the data over a short window. We chose 5 frames, given the time period of the motion changes in these cells. While the spot speed showed no discernible difference between tracks inside and those outside the boundary, both the windowed speed (Figure 3C; p-value <2.2x 10^-16^) and straightness (Figure 3D) were increased outside the spheroid boundary.

While easy to comprehend, the windowed speed is highly dependent on the window of both sampling and calculation. Furthermore, it does not in itself characterise the nature of the motion. An important descriptor that describes motion over a range of time-scales is the Mean Square Displacement (MSD). By plotting the MSD of a track at a range of time-scales, the motion of an object can be characterised (27). Our analysis suggests that, up to a time scale of several hours, cells in an untreated control spheroid are moving randomly (Figure 3G) with a diffusion coefficient of approximately 0.0055 *μ*m^2^/s. We further compared the diffusion coefficient of the track segments inside and outside of the spheroid (Figure 3H). While the two groups have similar random motion as indicated by the straight line in the MSD plot, the diffusion coefficient for inside tracks was slightly lower (0.0053 *μ*m^2^ versus 0.0068 *μ*m^2^ for outside tracks), corroborating our earlier findings with windowed speed.

### Analysis of radial components and track breakdown

As spheroids are spatially isotropic systems, the movement of cells is best described in spherical coordinates in relation to the centre of the spheroid. Velocity vectors can then be calculated for azimuthal, elevational and radial components (Figure 4A, B shows examples of the different coordinate systems for a 2D system). The radial velocity was plotted as a function of distance from the centre of the spheroid (Figure 4C) or as a function of time (Figure 4D). From our observations in the spheroid model, the sum of positive (i.e. outward) and negative (i.e. inward) velocities appear to approximate zero.

**Figure 4.**
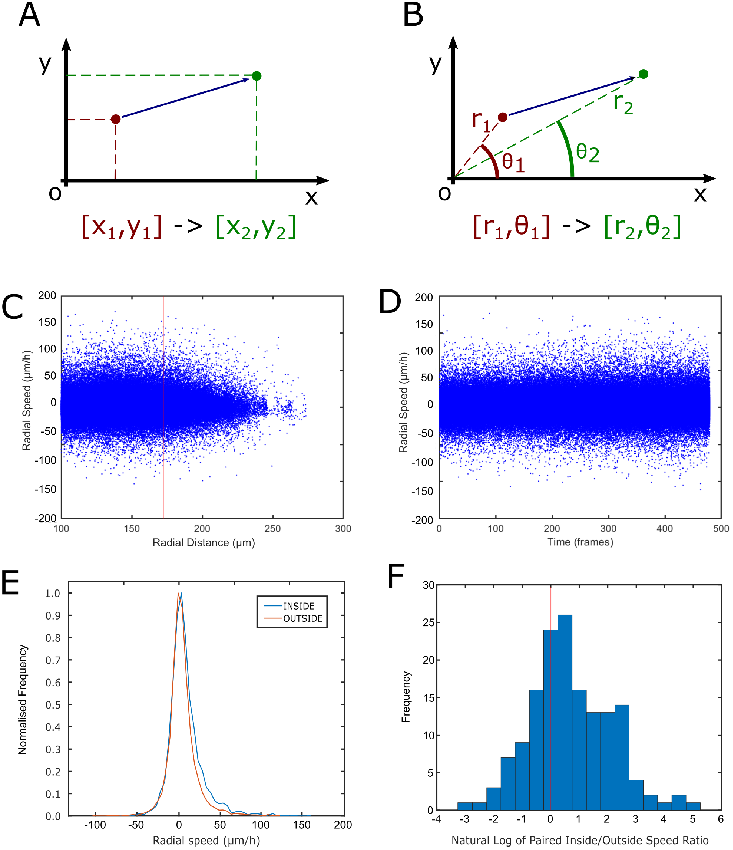
Analysis of radial components. The isotropic nature of a spheroid makes it well suited to a spherical coordinate system over Cartesian. (**A**) A 2D example of a Cartesian coordinate system and (**B**) of a circular coordinate system, which extends into 3D with the addition of ‘z’ and *ϕ* (phi) axes respectively. Radial velocity vectors are calculated and describe the movement in to (negative value) or out of (positive value) the spheroid. This can be investigated as a function of distance from the spheroid centroid (**C**) or time of the experiment (**D**). The red vertical line in (C) represents the calculated spheroid boundary (see Methods). Within the context of the spheroid boundary, the radial velocity vectors also allow interrogation of the difference in radial velocity (**E**) when moving within the spheroid (E: Inside) or in the surrounding matrix (E: Outside). Although not all cells leave the spheroid boundary, for cells that migrate (either way), the paired ratio (inside/outside parts of the same track) can be calculated to show relative changes in the radial velocity. The distribution of the natural log of the ratios is shown (**F**) where zero (red vertical line) represents no difference inside and out and a positive skew indicates slower speed inside. Panel (F) includes the results from three control experiments.

We wanted to investigate if migration patterns were different for cells migrating within spheroids compared with cells invading into the extra-spheroid space. The ‘migratory’ tracks (i.e. those which cross the boundary in either direction) were split into their inside and outside segments, taking into account multiple transitions. This allowed the comparison of all of the inside and outside components within an individual track. Despite the relatively few cells that migrate into the surrounding matrix over the course of the experiment (~15% of total number of cells detected), a slight positive skew is seen when the distribution of radial velocities is plotted separately for vectors inside and those outside the spheroid boundary (Figure 4E). Furthermore, by pairing the components of single tracks, it is possible to get a clearer picture of whether individual cells behave differently in the matrix compared to within the spheroid. The distribution of the natural log of the paired ratios are plotted in Figure 4F. The data are presented this way such that zero (shown with a red vertical line) represents no difference when comparing inside versus outside speeds. The positively-skewed data (with mean of the absolute values being 6.6) indicate the absolute radial velocity of cells decreases when migrating through the matrix compared with their time inside the spheroid.

### Testing anticancer drugs effects on cell migration and invasion

To assess the suitability of this 3D single cell-tracking approach as a method to evaluate inhibitors of invasion, we treated spheroids with 17-allylamino-17-demethoxygeldanamycin (17AAG) and cilengitide. Both drugs were added to the cell culture media during sample embedding in Matrigel. 17AAG has been shown to inhibit the growth and invasion of glioma cells by inhibiting heat shock protein 90 (HSP90), a chaperone protein that regulates numerous signalling proteins, including those involved in cell proliferation, survival, migration and invasion (28–30). Cilengitide is a cyclic peptide antagonist of integrins *α*v*β*3 and *α*v*β*5 that has recently been in clinical trials for recurrent and newly diagnosed glioblastoma (31). The evidence of the effect of cilengitide on invasion is mixed: while some reports document an anti-invasive effect (32, 33), others have found that it acts to promote cell detachment, migration and invasion in 2D cell-based assays (34). Similar to Maurer *et al*, we observed that cells treated with cilengitide showed more rounding up, presumably due to the impairment of adhesion properties (Supplemental Figure 2A). To clarify the role of integrins in the different steps involved in invasion, we initially tested the effects of cilengitide on 2D cell migration. Cilengitide slightly increased the distribution of mean track speeds compared to untreated controls and triggered cell detachment from the cell culture dish (Supplemental Figure 2B). However, when 3-day spheroids were treated with the same concentration of cilengitide, invasion into the matrix was not inhibited (Figure 5A, B and Supplemental Figure 3). The distribution of the windowed speed was comparable between control and cilengitide treated conditions (Figure 5F and Supplemental Movie 2). The absence of cilengitide effects in the 3D model conflicts with the detachment and changes in migration observed in 2D cell culture, and demonstrates the risks of extrapolating results observed in simple models to more complex ones, as the integrin composition is likely to change (35) (see discussion). Conversely, upon HSP90 inhibition with 17AAG, a lack of invasion was immediately evident from the time-lapse (see Figure 5D and Supplemental Movie 3). The cumulative distribution graph (Figure 5E and Supplemental Figure 3) was also markedly different from control (Figure 5A) and cilengitide treated spheroids (Figure 5B), with only a small amount of curve shift at the very edge of the spheroid, indicating severely restricted invasion into the matrix. The effects of 17AAG on cell invasion were independent of the cell-cycle inhibitory properties of this drug as, at the concentration used, we did not observe any effect on cell division over a 24 h time course in 2D cell culture (Supplemental Figure 4). The distribution of both the speed (Figure 5F) and straightness (Figure 5G) of cells in the 17AAG-treated spheroids were decreased compared with control and cilengitide treated spheroids, suggesting both slower and more tortuous motion overall.

**Figure 5.**
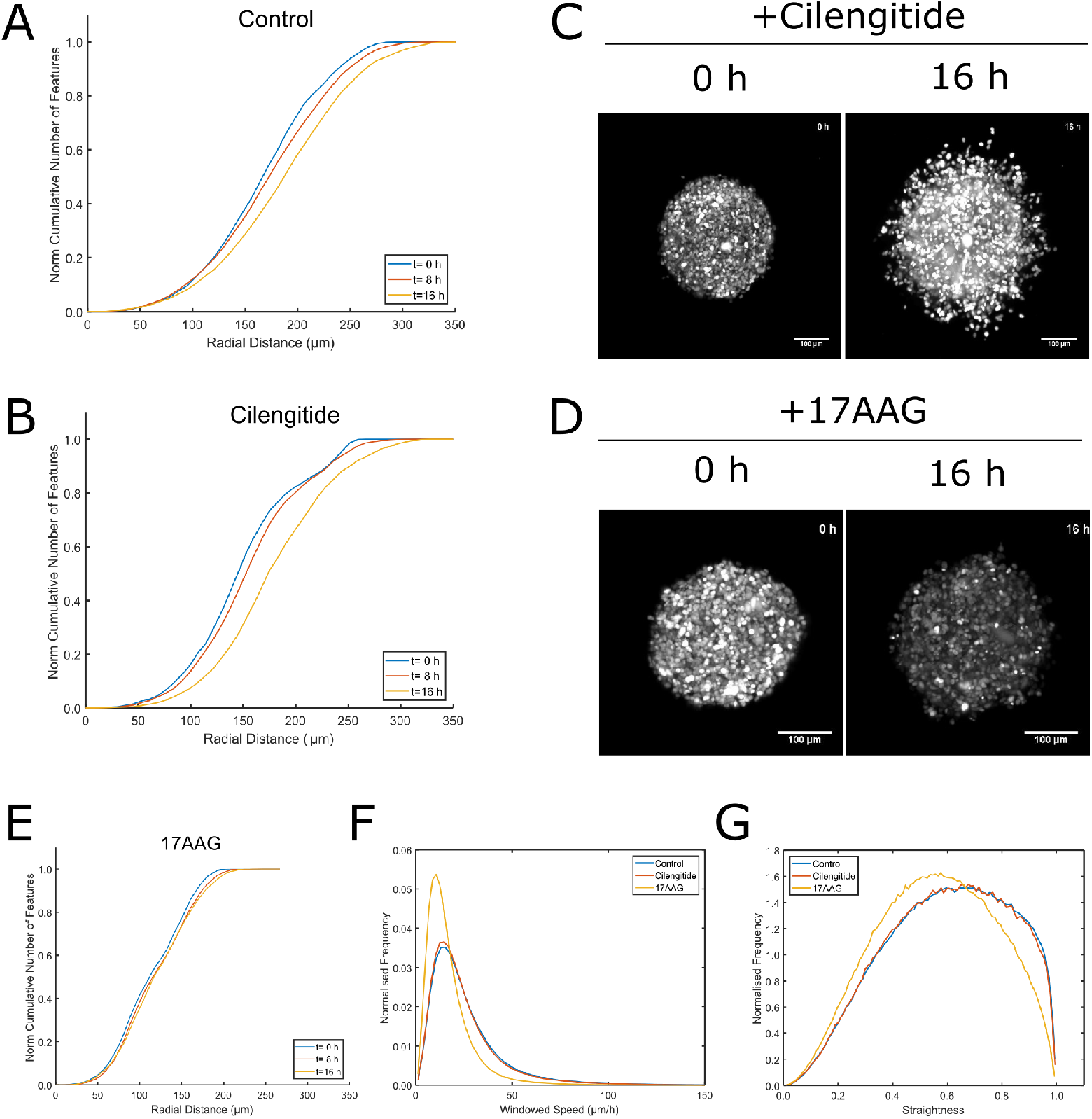
Perturbing migration with chemotherapeutics. The cumulative distribution of cells (as a function of distance from the spheroid centroid) in control (**A**) and cilengitide-treated (**B**) spheroids over time. (**C**) Maximum intensity projection for comparison of cilengitide spheroids with untreated control spheroids (see Figure 1A for comparison) at 0 h and 16 h. (**D**) Maximum intensity projection of 17AAG-treated spheroids invasion across the experimental time course. The cumulative distribution of cells (as a function of distance from the spheroid centroid) in 17AAG-treated spheroids (**E**) over time. (**F**) Distribution of the windowed speed. (**G**) Distribution of mean windowed straightness. Panels (A), (B), (E), (F) and (G) show the mean results from three experiments per condition. Panels (F) and (G) are normalised such that the area under the curve is the same.

In summary, we have developed a workflow (summarised in Supplemental Figure 1) for the analysis of spheroid cell migration and invasion data. The acquisition, postprocessing and parameter extraction of single cell movement in spheroids allows the effects of chemotherapeutic drugs on cell migration and invasion to be tested.

## Discussion

While spheroids have previously been used to investigate invasion, these investigations have until recently been limited by the available imaging techniques and a lack of clear metrics by which to quantify and compare conditions. Projecting 3D imaging data or relying on 2D imaging alone can make processing and analysis easier, yet this inherently decreases the accuracy of the measurements. We described here a workflow to aid in the acquisition and analysis of 4D data from lightsheet imaging, using spheroids derived from glioblastoma cells as a model system. By imaging and tracking individual cells, we were able to gain many of the broad insights provided in 2D acquisition systems as well as leveraging the detailed information gleaned from the 3D nature of the system.

### 3D cell microenvironment and migration speed

In line with *in vivo* observations, we recorded a high mobility of cells within gliobastoma spheroids. Under control conditions, the mean track speed of 27 *μ*m/h within the spheroid boundary correlates well with the results of Farin *et al.*, in which glioma cells injected into neonatal rat forebrains were tracked and evaluated to move at 25 *μ*m/h in sectioned tissues when imaged (as with our experiments) every 3 minutes (36). However, the tissue microenvironment and its properties are likely to affect cell migration. Indeed, we found that track speed and straightness were consistently increased in cells that were invading into the Matrigel compared with cells that were migrating within the spheroid (Figure 3D). It has been previously described how the speed and mode of cellular invasion are modulated by properties of the ECM such as composition, matrix stiffness, presence of growth factors and pore size (37–41). In addition, Jayatilaka *et al*. showed a remarkable dependence of cell speed on the distance between the cells in the matrix (42), reviewed by (43), with human breast carcinoma MDA-MB-231 cells migrating much more when they had close contacts. In our case, there were many more cell-to-cell contacts within the spheroid, which will have altered the migratory properties and ECM secreted by the cells. These parameters (secreted factors, cell-to cell contact and different ECM composition) can explain the different migration characteristics such as the straightness of the trajectories. Our experimental and analytical workflow provides a precise and quantitative measurement of the cell migration properties that can be used to assess these changes. One question that remains unanswered is whether cells migrating out from a spheroid are creating new pores, or using existing holes created by cells that have previously left. With slight adaptations such as using fluorescently labelled matrix or cell body markers, this workflow can be adapted to study the type of cell migration.

There are several advantages to nuclear tracking, chiefly that feature detection and segmentation during tracking is considerably more robust especially if cells are in contact. Touching cells expressing a soluble GFP or other cytosolic or membrane label are harder to segment within a complex 3D structure. Moreover, nuclear tracking allows the possibility of gathering additional information on cell division if required. However, light scattering reduced our ability to identify (or reliably track) features at the very centre of the spheroid. Additionally, some of the original images were acquired in single-view mode, therefore, the hemisphere furthest from the objective was obscured.

### Drug effects on cell invasion versus cell migration

To compare cell motion into the surrounding matrix to movement within the spheroid, a boundary has to be defined. The position of the boundary is somewhat arbitrary because there is a continuous transition in cell density between inside and outside of the spheroid. Here, we opted for a simple definition of ‘inside’ and ‘outside’ based on the initial time point. We found this to be a robust and useful functional definition that can be used to reliably segment tracks, as the ‘core’ of the spheroid does not appreciably change in size during the experiment. Of note, the open availability of the code makes it simple for others to adapt the boundary calculation or indeed any other part of the data processing.

Our results show that 17AAG prevented the invasion of cells into the Matrigel but interestingly did not prevent their migration within the spheroid, although the movement of cells inside 17AAG treated spheroids was slower and less directional than in control spheroids. 17AAG affects many signalling proteins through its inhibition of HSP90, making it difficult to pinpoint the mechanism through which invasion is inhibited (28, 29, 44). However, it has been shown that treatment with 17AAG and HSP90 inhibition modulates actin dynamics, thereby affecting cell migration and invasion (45).

In contrast to 17AAG, we found that cilengitide was not effective in inhibiting invasion and, furthermore, we could not detect any differences between spheroids treated with cilengitide and control spheroids. Cilengitide reduces cell viability by causing cell detachment (anoikis). The sensitivity of U87 cells to cilengitide-induced detachment has been shown to be dependent on the composition of the ECM, with cells cultured on collagen-coated plates resisting detachment (34, 46). Gritsenko and Friedl have recently demonstrated that *α*V and *β*1 integrins represent the primary adhesion systems for glioma cell migration in different migration models (spheroids and organotypic brain slices) (35). However, when they compared the same integrin interference strategy in the different models, they observed a variation in the contribution of integrin-dependent glioma invasion. Their results suggest that there are alternative cell-matrix interactions in complex brain-like models, which drive different cell migration properties and require specific inhibitors. Focal-adhesion- and integrin-independent cell migration, especially in 3D matrices, has been extensively reviewed by Paluch *et al*.(47). Specifically, the Sixt lab has shown, by ablating all integrins heterodimers from murine leucocytes, that integrin-independent migration still occurs in 3D environments(48). The discrepancy of the effects of integrin inhibition/ablation between 2D and 3D environments illustrates the importance of replicating environmental factors, such as ECM composition, to obtain a valid assessment of anti-invasive or antitumour activity and points to the fact that even more complex models than single cell type spheroids might be necessary for *in vitro* drug testing.

Nevertheless, the accurate measurement of cell velocity and movement, highly comparable to previous studies using glioma cells by Farin *et al*. (36), supports the use of live spheroid imaging in drug screening prior to costly investigations in preclinical animal models. Moreover, our approach has overcome some of the limitations faced by Farin *et al.*, who were only able to track the movements of cells in the x-y plane(36). In conclusion, light sheet microscopy provides an excellent method for measuring cellular movements in multicellular tumour spheroids, providing not only high quality time-lapse images but also a framework to track and analyse these dense structures. Our findings with the two anti-cancer drugs tested indicate that it also holds promise for screening and investigating the mechanism of anti-invasive drugs, contributing to the principles of the replacement and reduction of the use of animals in experimental research.

## ACKNOWLEDGEMENTS

The microscopes used for this work were funded in part by MRC grant MR/K015931/1 and BBSRC Alert13 grant BB/L014947/1. RR was funded by Naseem’s Manx Brain Tumour Charity. This work is also supported by EPSRC grant EP/N014499/1 ‘EPSRC Centre for New Mathematical Sciences Capabilities for Healthcare Technologies’. We would like to thank Marco Marcello and the Centre for Cell Imaging for provision of training, access to equipment and experimental support.

## AUTHOR CONTRIBUTIONS

RR designed the experimental protocols, performed all molecular biology, cell culture and microscopy experiments and analysed the data. DM wrote the code for data processing and implemented the image analysis workflow. DM contributed to image data analysis and interpretation. RB and RL provided help and ideas for mathematical interpretation of the data, statistical analysis and for choosing the most relevant output parameters. RK performed the experiment with fluorescently labelled matrix. VS conceived and coordinated the study. RR, DM and VS wrote the manuscript. DM prepared all the figures. All authors reviewed drafts of the manuscript and gave final approval.

## Supplementary Data

**Figure S1.**
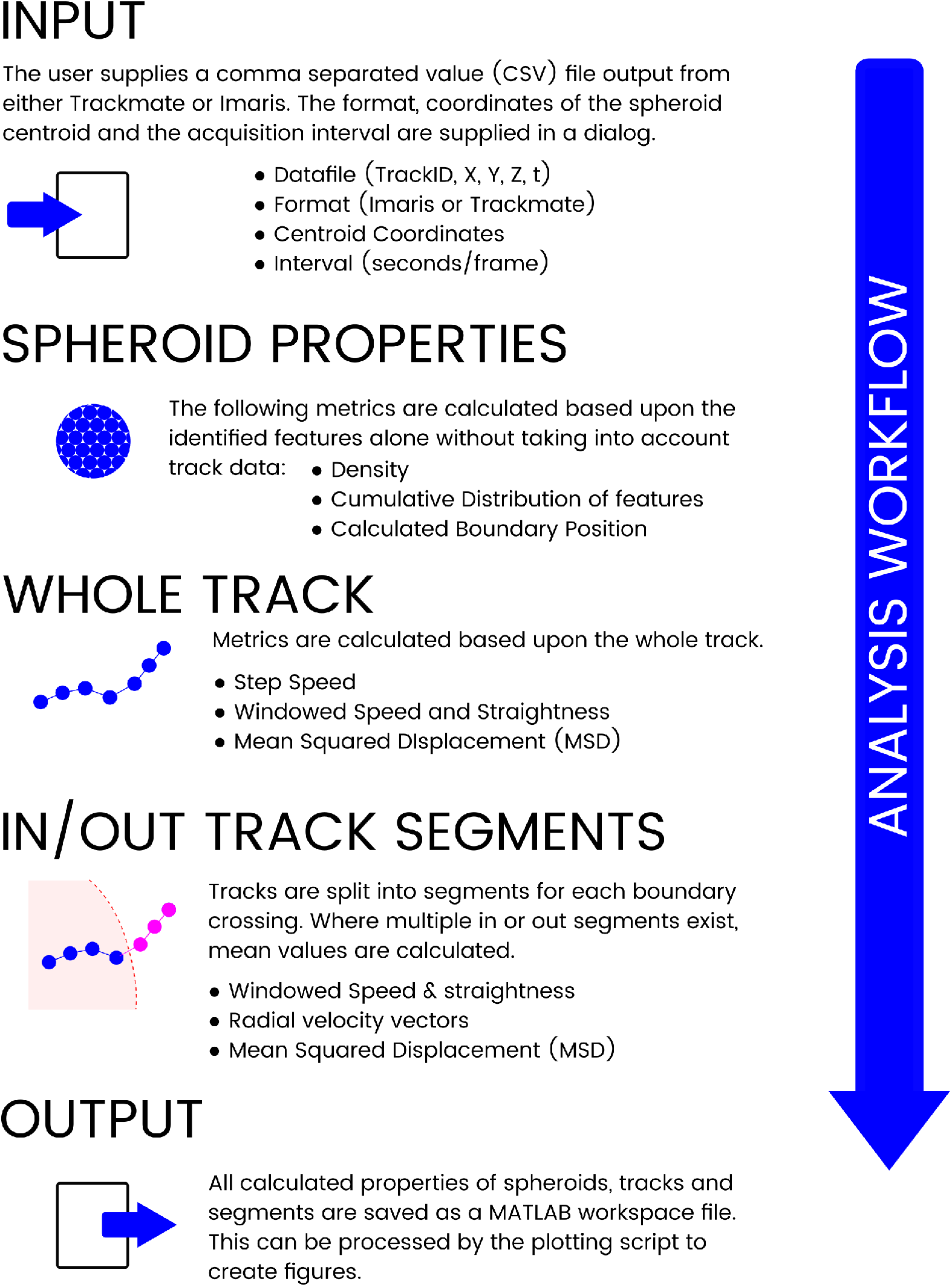
Main steps in the track analysis workflow. The main inputs, outputs and processing steps are briefly highlighted. Data are analysed based on bulk features to characterise the motion of the whole spheroid, whole track data and on segments, which are defined as parts of a track existing inside or outside the spheroid boundary. More detail can be found in the heavily-commented code (see Code Availability in Materials and Methods).

**Figure S2.**
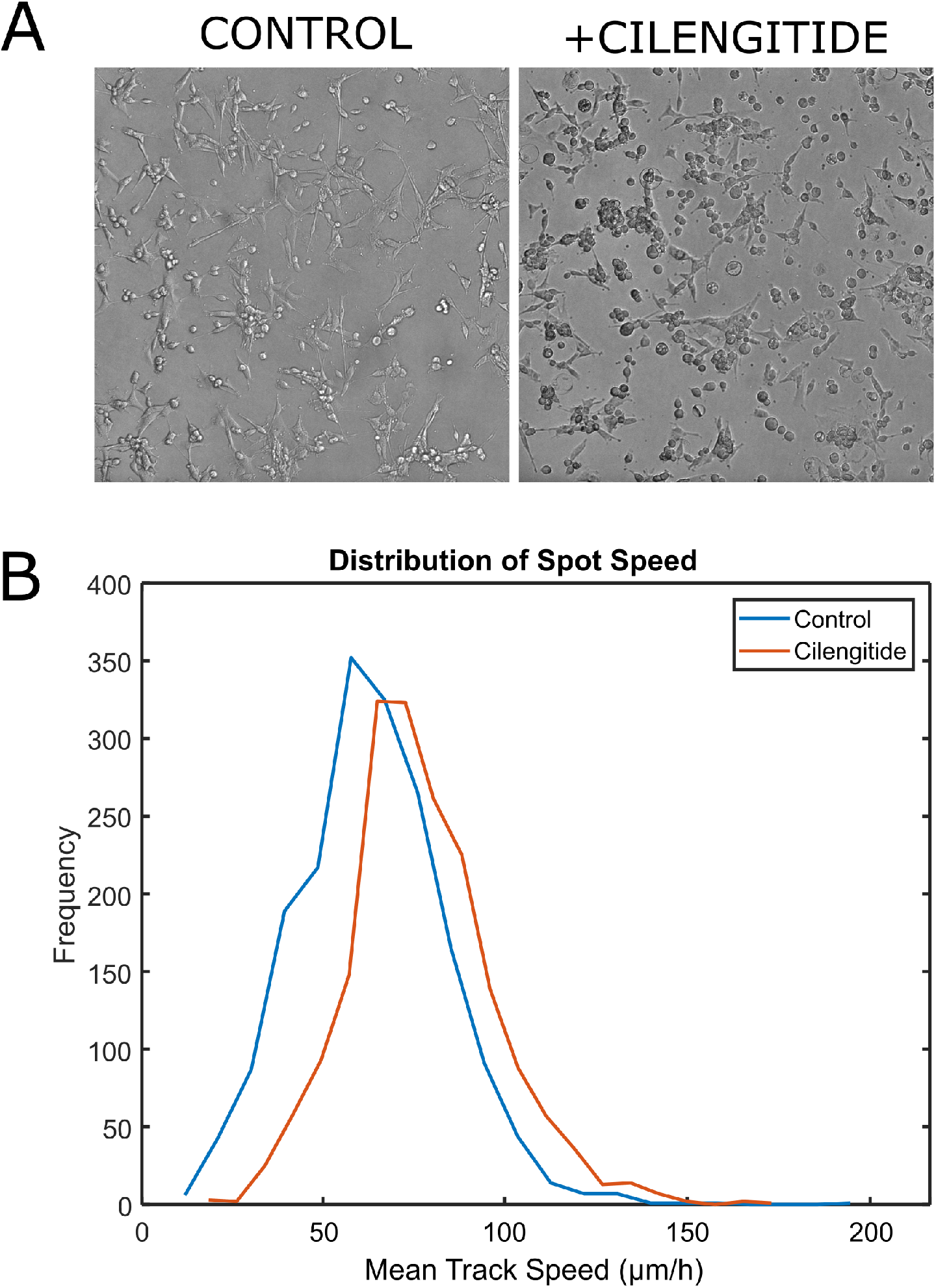
(**A**) Cilengitide treatment in 2D culture. U87MG glioblastoma cells grown in 2D cell culture show a more rounded, less adhesive phenotype when pre-treated with cilengitide. (**B**) When tracked over 16 hours, cilengitide-treated cells show a slightly faster track speed.

**Figure S3.**
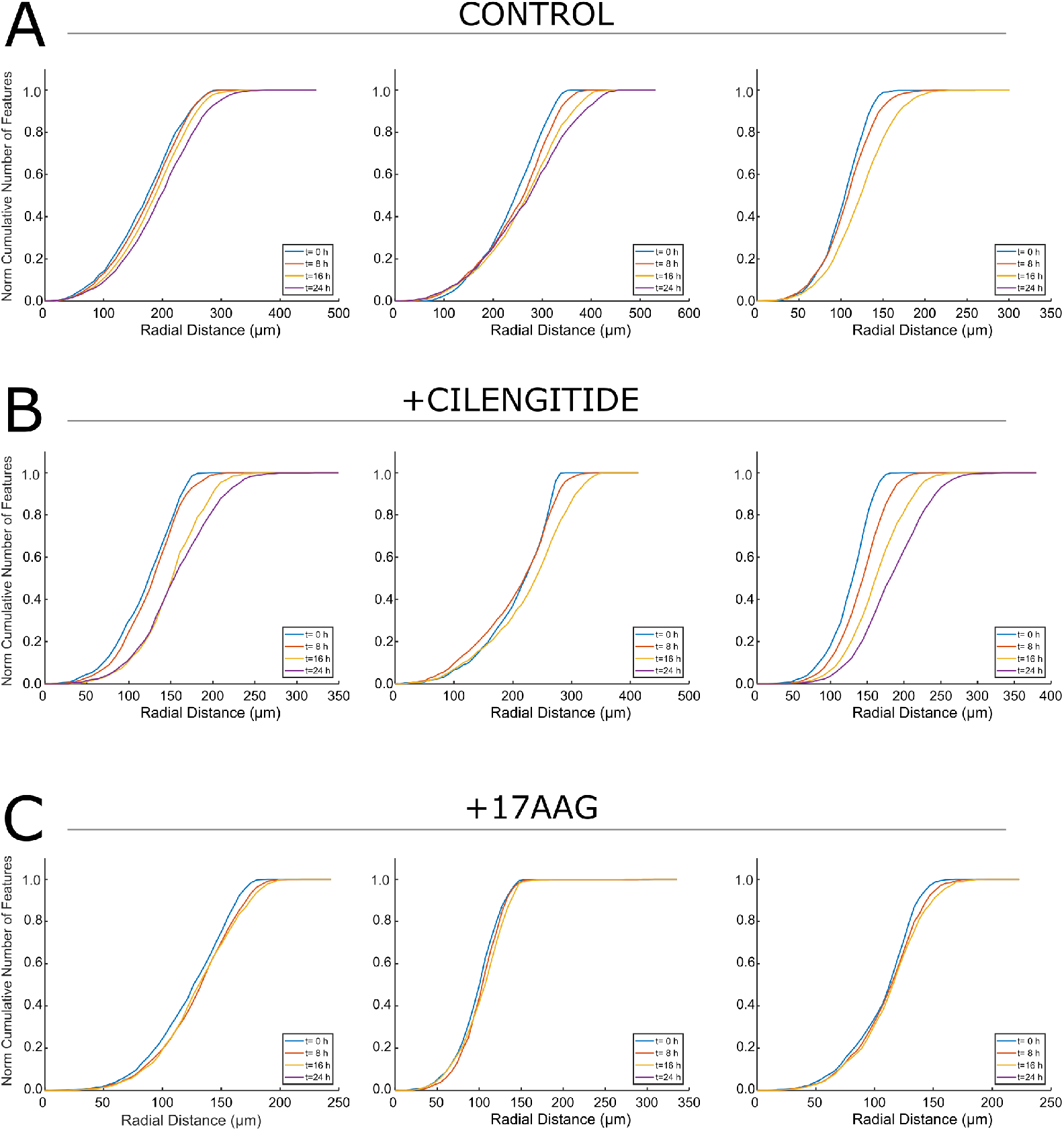
The cumulative distribution of cells (as a function of distance from the spheroid centroid) is shown for three replicates of control (**A**), cilengitide-treated (**B**) and 17AAG-treated (**C**) spheroids over time.

**Figure S4.**
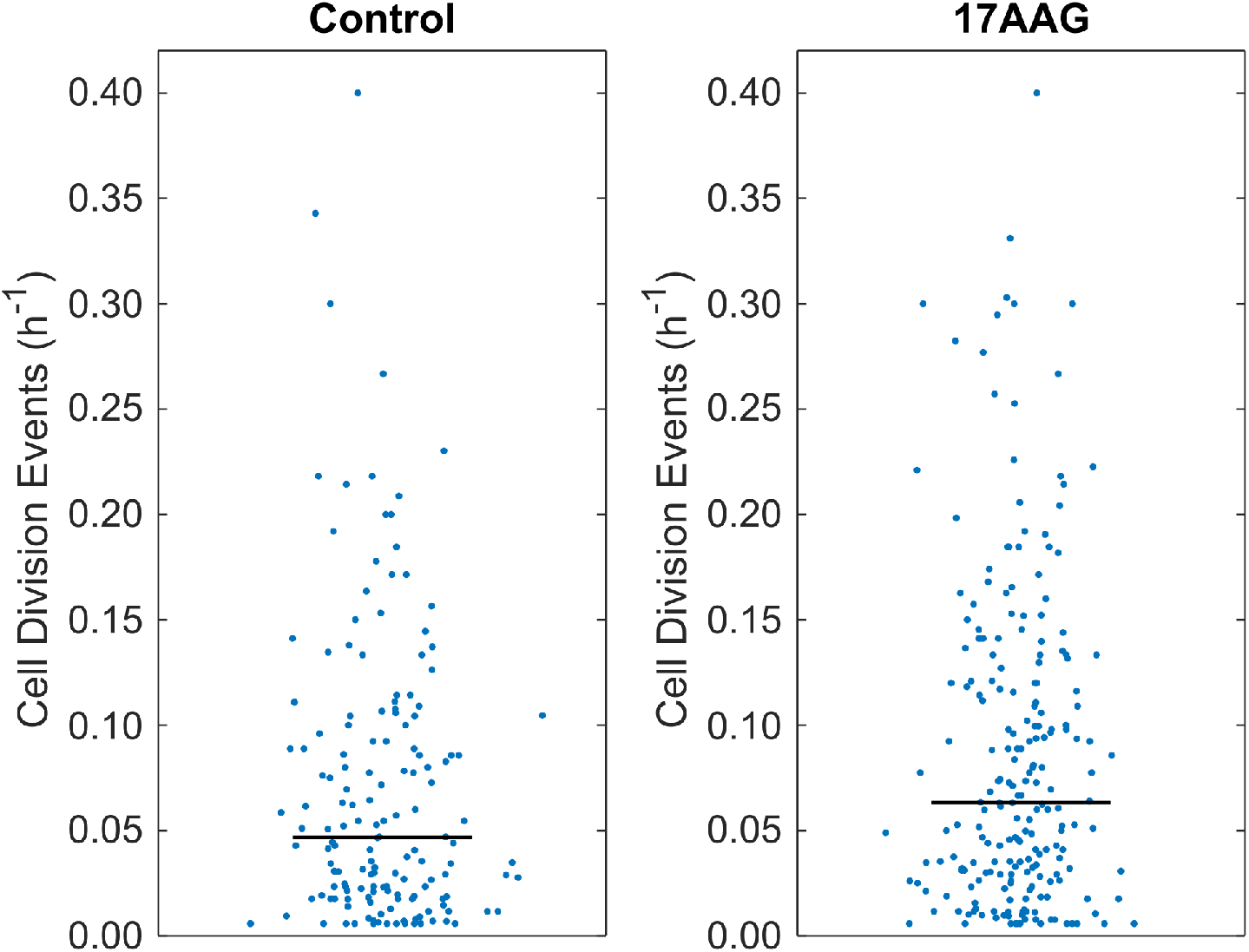
The number of “Track Split” events (used as a proxy for cell division) were recorded for untreated and 17AAG-treated U87 H2B-mRFP cells grown in 2D culture. At least 150 tracks were analysed for each plot.

**Figure S5.**
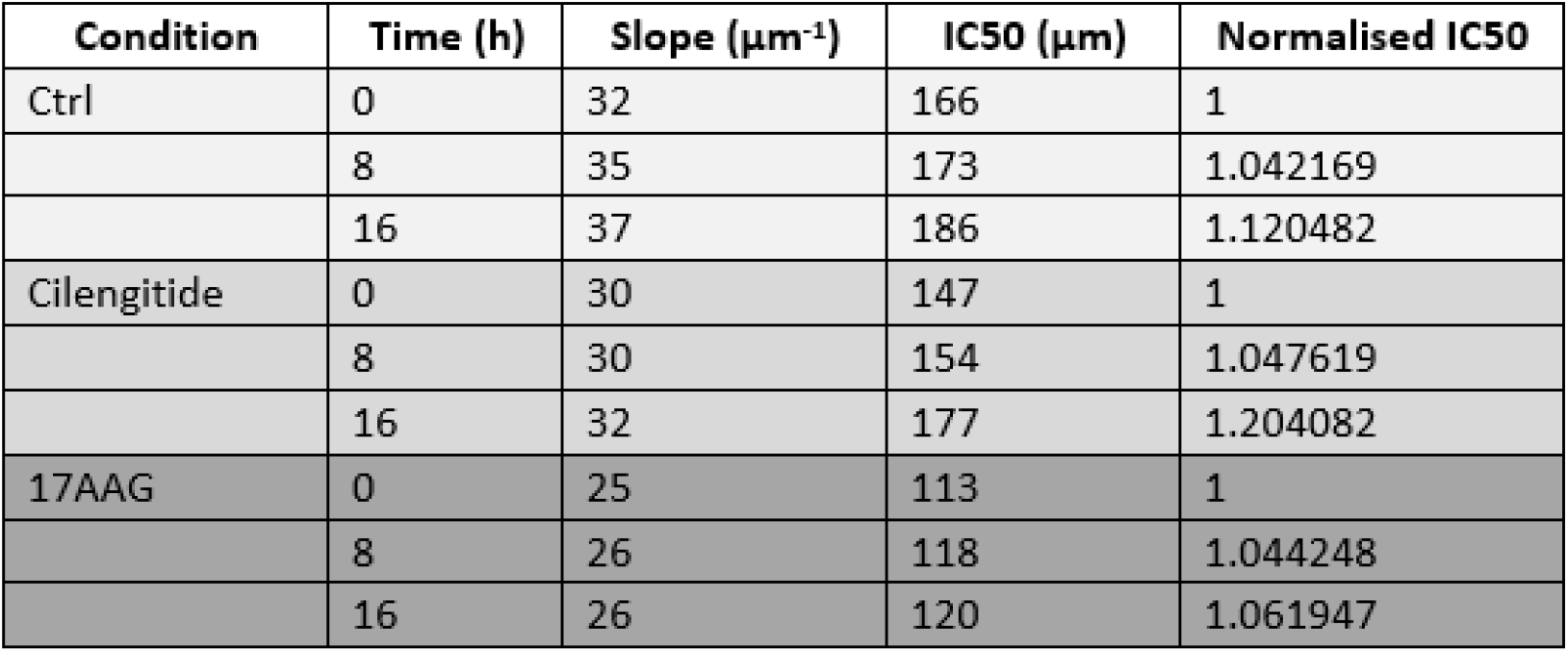
Fit parameters calculated for Cumulative Distribution plots for each experimental condition

## Supplementary Movies

**Supplementary Movie 1.** Invasion into the matrix. Glioblastoma spheroids made up of U87 cells expressing H2B-RFP were grown for 3 days, mounted on a lightsheet microscope (0 h) and imaged for up to 24 h. The movie is rendered using Imaris.

**Supplementary Movie 2.** Cilengitide treatment does not affect invasion. Glioblastoma spheroids made up of U87 cells expressing H2B-RFP were grown for 3 days, mounted on a lightsheet microscope with 2 *μ*M cilengitide (0 h) and imaged for up to 24 h. The movie shows a maximum intensity projection through the z-plane.

**Supplementary Movie 3.** 17AAG treatment affect both migration and invasion. Glioblastoma spheroids made up of U87 cells expressing H2B-RFP were grown for 3 days, mounted on a lightsheet microscope with 0.5 *μ*M 17AAG (0 h) and imaged for up to 24 h. The movie shows a maximum intensity projection through the z-plane.

## Bibliography

1. S. Walenta, J. Doetsch, W. Mueller-Klieser, and L. A. Kunz-Schughart. Metabolic imaging in multicellular spheroids of oncogene-transfected fibroblasts. J Histochem Cytochem, 48 (4):509–22, 2000. ISSN 0022-1554 (Print) 0022-1554 (Linking).

2. Jennifer Laurent, Celine Frongia, Martine Cazales, Odile Mondesert, Bernard Ducommun, and Valerie Lobjois. Multicellular tumor spheroid models to explore cell cycle checkpoints in 3d. Bmc Cancer, 13, 2013. ISSN 1471-2407. doi: 10.1186/1471-2407-13-73.

3. Pawel Swietach, Shalini Patiar, Claudiu T. Supuran, Adrian L. Harris, and Richard D. Vaughan-Jones. The role of carbonic anhydrase 9 in regulating extracellular and intracellular ph in three-dimensional tumor cell growths. Journal of Biological Chemistry, 284 (30):20299–20310, 2009. ISSN 0021-9258. doi: 10.1074/jbc.M109.006478.

4. D. Hanahan and R. A. Weinberg. The hallmarks of cancer. Cell, 100(1):57–70, 2000. ISSN 0092-8674 (Print) 0092-8674 (Linking).

5. Nina Kramer, Angelika Walzl, Christine Unger, Margit Rosner, Georg Krupitza, Markus Hengstschlaeger, and Helmut Dolznig. In vitro cell migration and invasion assays. Mutation Research-Reviews in Mutation Research, 752(1):10–24, 2013. ISSN 1383-5742. doi: 10.1016/j.mrrev.2012.08.001.

6. Brendon M. Baker and Christopher S. Chen. Deconstructing the third dimension - how 3d culture microenvironments alter cellular cues. Journal of Cell Science, 125(13):3015–3024, 2012. ISSN 0021-9533. doi: 10.1242/jcs.079509.

7. S. Breslin and L. O’Driscoll. Three-dimensional cell culture: the missing link in drug discovery. Drug Discov Today, 18(5-6):240–9, 2013. ISSN 1878-5832 (Electronic) 1359-6446 (Linking). doi: 10.1016/j.drudis.2012.10.003.

8. M. Vinci, S. Gowan, F. Boxall, L. Patterson, M. Zimmermann, W. Court, C. Lomas, M. Mendiola, D. Hardisson, and S. A. Eccles. Advances in establishment and analysis of threedimensional tumor spheroid-based functional assays for target validation and drug evaluation. Bmc Biology, 10, 2012. ISSN 1741-7007. doi: Artn2910.1186/1741-7007-10-29.

9. Lin Cheng, Qiulian Wu, Olga A. Guryanova, Zhi Huang, Qian Huang, Jeremy N. Rich, and Shideng Bao. Elevated invasive potential of glioblastoma stem cells. Biochemical and Biophysical Research Communications, 406(4), 2011. ISSN 0006-291X. doi: 10.1016/j.bbrc.2011.02.123.

10. Andrew M. Stein, Tim Demuth, David Mobley, Michael Berens, and Leonard M. Sander. A mathematical model of glioblastoma tumor spheroid invasion in a three-dimensional in vitro experiment. Biophysical Journal, 92(1):356–365, 2007. ISSN 0006-3495. doi: 10.1529/biophysj.106.093468.

11. S. Blacher, C. Erpicum, B. Lenoir, J. Paupert, G. Moraes, S. Ormenese, E. Bullinger, and A. Noel. Cell invasion in the spheroid sprouting assay: A spatial organisation analysis adaptable to cell behaviour. Plos One, 9(5), 2014. ISSN 1932-6203. doi: ARTNe9701910.1371/journal.pone.0097019.

12. E. G. Reynaud, U. Krzic, K. Greger, and E. H. Stelzer. Light sheet-based fluorescence microscopy: more dimensions, more photons, and less photodamage. HFSP J, 2(5):266–75, 2008. ISSN 1955-2068 (Print) 1955-205X (Linking). doi: 10.2976/1.2974980.

13. E. H. Stelzer. Light-sheet fluorescence microscopy for quantitative biology. Nat Methods, 12 (1):23–6, 2015. ISSN 1548-7105 (Electronic) 1548-7091 (Linking). doi: 10.1038/nmeth.3219.

14. J. Huisken, J. Swoger, F. Del Bene, J. Wittbrodt, and E. H. Stelzer. Optical sectioning deep inside live embryos by selective plane illumination microscopy. Science, 305(5686):1007–9, 2004. ISSN 1095-9203 (Electronic) 0036-8075 (Linking). doi: 10.1126/science.1100035.

15. Y. Hou, J. Konen, D. J. Brat, A. I. Marcus, and L. A. D. Cooper. Tasi: A software tool for spatial-temporal quantification of tumor spheroid dynamics. Sci Rep, 8(1):7248, 2018. ISSN 2045-2322 (Electronic) 2045-2322 (Linking). doi: 10.1038/s41598-018-25337-4.

16. C. Lorenzo, C. Frongia, R. Jorand, J. Fehrenbach, P. Weiss, A. Maandhui, G. Gay, B. Ducommun, and V. Lobjois. Live cell division dynamics monitoring in 3d large spheroid tumor models using light sheet microscopy. Cell Div, 6:22, 2011. ISSN 1747-1028 (Electronic) 1747-1028 (Linking). doi: 10.1186/1747-1028-6-22.

17. A. Desmaison, L. Guillaume, S. Triclin, P. Weiss, B. Ducommun, and V. Lobjois. Impact of physical confinement on nuclei geometry and cell division dynamics in 3d spheroids. Sci Rep, 8(1):8785, 2018. ISSN 2045-2322 (Electronic) 2045-2322 (Linking). doi: 10.1038/s41598-018-27060-6.

18. Kazuhiko Kurozumi, Tomotsugu Ichikawa, Manabu Onishi, Kentaro Fujii, and Isao Date. Cilengitide treatment for malignant glioma: Current status and future direction. Neurologia Medico-Chirurgica, 52(8):539–547, 2012. ISSN 0470-8105.

19. Robert H. Kutner, Xian-Yang Zhang, and Jakob Reiser. Production, concentration and titration of pseudotyped hiv-1-based lentiviral vectors. Nature Protocols, 4(4):495–505, 2009. ISSN 1754-2189. doi: 10.1038/nprot.2009.22.

20. M. Marcello, R. Richards, D. Mason, and V. See. Live imaging of cell invasion using a multicellular spheroid model and light-sheet microscopy. Adv Exp Med Biol, 1035:155–161, 2017. ISSN 0065-2598 (Print) 0065-2598 (Linking). doi: 10.1007/978-3-319-67358-5_11.

21. J. Schindelin, I. Arganda-Carreras, E. Frise, V. Kaynig, M. Longair, T. Pietzsch, S. Preibisch, C. Rueden, S. Saalfeld, B. Schmid, J. Y. Tinevez, D. J. White, V. Hartenstein, K. Eliceiri, P. Tomancak, and A. Cardona. Fiji: an open-source platform for biological-image analysis. Nat Methods, 9(7):676–82, 2012. ISSN 1548-7105 (Electronic) 1548-7091 (Linking). doi: 10.1038/nmeth.2019.

22. S. Preibisch, S. Saalfeld, J. Schindelin, and P. Tomancak. Software for bead-based registration of selective plane illumination microscopy data. Nat Methods, 7(6):418–9, 2010. ISSN 1548-7105 (Electronic) 1548-7091 (Linking). doi: 10.1038/nmeth0610-418.

23. N. Chenouard, I. Smal, F. de Chaumont, M. Maska, I. F. Sbalzarini, Y. Gong, J. Cardinale, C. Carthel, S. Coraluppi, M. Winter, A. R. Cohen, W. J. Godinez, K. Rohr, Y. Kalaidzidis, L. Liang, J. Duncan, H. Shen, Y. Xu, K. E. Magnusson, J. Jalden, H. M. Blau, P. Paul-Gilloteaux, P. Roudot, C. Kervrann, F. Waharte, J. Y. Tinevez, S. L. Shorte, J. Willemse, K. Celler, G. P. van Wezel, H. W. Dan, Y. S. Tsai, C. Ortiz de Solorzano, J. C. Olivo-Marin, and E. Meijering. Objective comparison of particle tracking methods. Nat Methods, 11(3): 281-9, 2014. ISSN 1548-7105 (Electronic) 1548-7091 (Linking). doi: 10.1038/nmeth.2808.

24. J. Y. Tinevez, N. Perry, J. Schindelin, G. M. Hoopes, G. D. Reynolds, E. Laplantine, S. Y. Bednarek, S. L. Shorte, and K. W. Eliceiri. Trackmate: An open and extensible platform for single-particle tracking. Methods, 115:80–90, 2017. ISSN 1095-9130 (Electronic) 1046-2023 (Linking). doi: 10.1016/j.ymeth.2016.09.016.

25. M. Vinci, C. Box, M. Zimmermann, and S. A. Eccles. Tumor spheroid-based migration assays for evaluation of therapeutic agents. Methods Mol Biol, 986:253–66, 2013. ISSN 1940-6029 (Electronic) 1064-3745 (Linking). doi: 10.1007/978-1-62703-311-4_16.

26. D.B. Schott, M. Traub, C. Schlagenhauf, M. Takamiya, T. Antritter, A. Bartschat, K. Loffler, Blessing, J. C. Otte, A. Y. Kobitski, G. U. Nienhaus, U. Strahle, R. Mikut, and J. Stegmaier. Embryominer: A new framework for interactive knowledge discovery in large-scale cell tracking data of developing embryos. PLoS Comput Biol, 14(4):e1006128, 2018. ISSN 1553-7358 (Electronic) 1553-734X (Linking). doi: 10.1371/journal.pcbi.1006128.

27. A. Kusumi, Y. Sako, and M. Yamamoto. Confined lateral diffusion of membrane receptors as studied by single particle tracking (nanovid microscopy). effects of calcium-induced differentiation in cultured epithelial cells. Biophys J, 65(5):2021–40, 1993. ISSN 0006-3495 (Print) 0006-3495 (Linking). doi: 10.1016/S0006-3495(93)81253-0.

28. Shinji Tsutsumi, Kristin Beebe, and Len Neckers. Impact of heat-shock protein 90 on cancer metastasis. Future Oncology, 5(5):679–688, 2009. ISSN 1479-6694. doi: 10.2217/fon.09.30.

29. Claire Marie-Elisabeth Sauvageot, Jessica Leigh Weatherbee, Santosh Kesari, Susan Elizabeth Winters, Jessica Barnes, Jamie Dellagatta, Naren Raj Ramakrishna, Charles Dean Stiles, Andrew Li-Jen Kung, Mark W. Kieran, and Patrick Yung Chih Wen. Efficacy of the hsp90 inhibitor 17-aag in human glioma cell lines and tumorigenic glioma stem cells. Neuro Oncology, 11(2):109–121, 2009. ISSN 1522-8517. doi: 10.1215/15228517-2008-060.

30. K. Miekus, J. Kijowski, M. Sekula, and M. Majka. 17aep-ga, an hsp90 antagonist, is a potent inhibitor of glioblastoma cell proliferation, survival, migration and invasion. Oncol Rep, 28 (5):1903–9, 2012. ISSN 1791-2431 (Electronic) 1021-335X (Linking). doi: 10.3892/or.2012.1996.

31. O. L. Chinot. Cilengitide in glioblastoma: when did it fail? Lancet Oncol, 15(10):1044–5, 2014. ISSN 1474-5488 (Electronic) 1470-2045 (Linking). doi: 10.1016/S1470-2045(14)70403-6.

32. Manabu Onishi, Tomotsugu Ichikawa, Kazuhiko Kurozumi, Kentaro Fujii, Koichi Yoshida, Satoshi Inoue, Hiroyuki Michiue, E. Antonio Chiocca, Balveen Kaur, and Isao Date. Bimodal anti-glioma mechanisms of cilengitide demonstrated by novel invasive glioma models. Neuropathology, 33(2):162–174, 2013. ISSN 0919-6544. doi: 10.1111/j.1440-1789.2012.01344.x.

33. A. Wilisch-Neumann, N. Kliese, D. Pachow, T. Schneider, J. P. Warnke, W. E. Braunsdorf, F. D. Bohmer, P. Hass, D. Pasemann, C. Helbing, E. Kirches, and C. Mawrin. The integrin inhibitor cilengitide affects meningioma cell motility and invasion. Clin Cancer Res, 19(19): 5402–12, 2013. ISSN 1078-0432 (Print) 1078-0432 (Linking). doi: 10.1158/1078-0432.CCR-12-0299.

34. G. D. Maurer, I. Tritschler, B. Adams, G. Tabatabai, W. Wick, R. Stupp, and M. Weller. Cilengitide modulates attachment and viability of human glioma cells, but not sensitivity to irradiation or temozolomide in vitro. Neuro Oncol, 11(6):747–56, 2009. ISSN 1523-5866. doi: 10.1215/15228517-2009-012.

35. P. G. Gritsenko and P. Friedl. Adaptive adhesion systems mediate glioma cell invasion in complex environments. Journal of Cell Science, 131(15), 2018. ISSN 0021-9533. doi: UNSPjcs21638210.1242/jcs.216382.

36. A. Farin, S. O. Suzuki, M. Weiker, J. E. Goldman, J. N. Ruce, and P. Canoll. Transplanted glioma cells migrate and proliferate on host brain vasculature: A dynamic analysis. Glia, 53 (8):799–808, 2006. ISSN 0894-1491. doi: 10.1002/glia.20334.

37. R. H. Goldbrunner, H. K. Haugland, C. E. Klein, S. Kerkau, K. Roosen, and J. C. Tonn. Ecm dependent and integrin mediated tumor cell migration of human glioma and melanoma cell lines under serum-free conditions. Anticancer Research, 16(6B):3679–3687, 1996. ISSN 0250-7005.

38. Marisol Herrera-Perez, Sherry L. Voytik-Harbin, and Jenna L. Rickus. Extracellular matrix properties regulate the migratory response of glioblastoma stem cells in three-dimensional culture. Tissue Engineering Part A, 21(19-20):2572–2582, 2015. ISSN 1937-3341. doi: 10.1089/ten.tea.2014.0504.

39. K. Wolf, M. te Lindert, M. Krause, S. Alexander, J. te Riet, A. L. Willis, R. M. Hoffman, C. G. Figdor, S. J. Weiss, and P. Friedl. Physical limits of cell migration: Control by ecm space and nuclear deformation and tuning by proteolysis and traction force. Journal of Cell Biology, 201(7):1069–1084, 2013. ISSN 0021-9525. doi: 10.1083/jcb.201210152.

40. Muhammad H. Zaman, Linda M. Trapani, Alisha Siemeski, Drew MacKellar, Haiyan Gong, Roger D. Kamm, Alan Wells, Douglas A. Lauffenburger, and Paul Matsudaira. Migration of tumor cells in 3d matrices is governed by matrix stiffness along with cell-matrix adhesion and proteolysis. Proceedings of the National Academy of Sciences of the United States of America, 103(29):10889–10894, 2006. ISSN 0027-8424. doi: 10.1073/pnas.0604460103.

41. M. F. Ware, A. Wells, and D. A. Lauffenburger. Epidermal growth factor alters fibroblast migration speed and directional persistence reciprocally and in a matrix-dependent manner. J Cell Sci, 111 (Pt 16):2423–32, 1998. ISSN 0021-9533 (Print) 0021-9533 (Linking).

42. H. Jayatilaka, P. Tyle, J. J. Chen, M. Kwak, J. Ju, H. J. Kim, J. S. H. Lee, P. H. Wu, D. M. Gilkes, R. Fan, and D. Wirtz. Synergistic il-6 and il-8 paracrine signalling pathway infers a strategy to inhibit tumour cell migration. Nat Commun, 8:15584, 2017. ISSN 2041-1723 (Electronic) 2041-1723 (Linking). doi: 10.1038/ncomms15584.

43. P. H. Wu, D. M. Gilkes, and D. Wirtz. The biophysics of 3d cell migration. Annual Review of Biophysics, Vol 47, 47:549–567, 2018. ISSN 1936-122x. doi: 10.1146/annurev-biophys-070816-033854.

44. W. B. Pratt and D. O. Toft. Regulation of signaling protein function and trafficking by the hsp90/hsp70-based chaperone machinery. Experimental Biology and Medicine, 228(2): 111–133, 2003. ISSN 1535-3702.

45. A. Taiyab and M. Rao Ch. Hsp90 modulates actin dynamics: inhibition of hsp90 leads to decreased cell motility and impairs invasion. Biochim Biophys Acta, 1813(1):213–21, 2011. ISSN 0006-3002 (Print) 0006-3002 (Linking). doi: 10.1016/j.bbamcr.2010.09.012.

46. P. Leblond, A. Dewitte, F. Le Tinier, C. Bal-Mahieu, M. Baroncini, T. Sarrazin, E. Lartigau, A. Lansiaux, and S. Meignan. Cilengitide targets pediatric glioma and neuroblastoma cells through cell detachment and anoikis induction. Anticancer Drugs, 24(8):818–25, 2013. ISSN 1473-5741. doi: 10.1097/CAD.0b013e328362edc5.

47. E. K. Paluch, I. M. Aspalter, and M. Sixt. Focal adhesion-independent cell migration. Annu Rev Cell Dev Biol, 32:469–490, 2016. ISSN 1530-8995 (Electronic) 1081-0706 (Linking). doi: 10.1146/annurev-cellbio-111315-125341.

48. T. Lammermann, B. L. Bader, S. J. Monkley, T. Worbs, R. Wedlich-Soldner, K. Hirsch, M. Keller, R. Forster, D. R. Critchley, R. Fassler, and M. Sixt. Rapid leukocyte migration by integrin-independent flowing and squeezing. Nature, 453(7191):51–5, 2008. ISSN 1476-4687 (Electronic) 0028-0836 (Linking). doi: 10.1038/nature06887

